## Supplementary Table 3 for "Structural Insights into The Role of MAGEA4 in RAD18 Regulation: Implications for Ubiquitin Ligase-Binding across the MAGE Protein Family"

**Supplementary table 3:** Ubiquitination sites of Rad18 and ubiquitin chain linkages detected in the autoubiquitinated Rad18/Rad6 sample

|  | Modified residue | Number of reads | diGly Score |
| --- | --- | --- | --- |
| Rad18  Residue | K83 | 2 | 45,4 |
|  | K102 | 4 | 77,8 |
|  | K110 | 5 | 98,9 |
|  | K115 | 4 | 81,1 |
|  | K151 | 6 | 68,5 |
|  | K154 | 20 | 69,1 |
|  | K156 | 22 | 77,4 |
|  | K161 | 9 | 63,6 |
|  | K168 | 8 | 68,5 |
|  | K170 | 3 | 60,7 |
|  | K186 | 5 | 64,1 |
|  | K197 | 4 | 66,9 |
|  | K241 | 0 | N/A |
|  | K245 | 3 | 51,0 |
|  | K257 | 2 | 50,7 |
|  | K258 | 1 | 52,4 |
|  | K259 | 1 | 71,0 |
|  | K261 | 8 | 60,3 |
|  | K271 | 11 | 67,8 |
|  | K276 | 6 | 72,2 |
|  | K309 | 4 | 73,1 |
|  | K318 | 24 | 63,3 |
|  | K328 | 17 | 77,2 |
|  | K333 | 5 | 87,8 |
|  | K341 | 6 | 88,5 |
|  | K345 | 2 | 76,5 |
|  | K347 | 4 | 65,5 |
|  | K363 | 2 | 46,0 |
|  | K370 | 14 | 66,0 |
|  | K376 | 6 | 74,8 |
|  | K383 | 2 | 80,6 |
|  | K462 | 9 | 150,4 |
| Ubiquitin  Residue | K63 | 24 | 83,1 |
|  | K48 | 15 | 56,9 |
|  | K11 | 16 | 66,5 |
|  | K6 | 12 | 35,8 |
